# Hematopoietic Stem and Progenitor Cells Use Podosomes to Transcellularly Home to the Bone Marrow

**DOI:** 10.1101/269704

**Authors:** Timo Rademakers, Marieke Goedhart, Mark Hoogenboezem, Alexander García Ponce, Jos van Rijssel, Maryna Samus, Michael Schnoor, Stefan Butz, Stephan Huveneers, Dietmar Vestweber, Martijn A. Nolte, Carlijn Voermans, Jaap D. van Buul

**Affiliations:** Dept. Plasma Proteins, Sanquin Research and Landsteiner Laboratory, Academic Medical Center, Amsterdam, the Netherlands; Dept. Hematopoiesis, Sanquin Research and Landsteiner Laboratory, Academic Medical Center, Amsterdam, the Netherlands; Department for Molecular Biomedicine, Center of Research and Advanced Studies (CINVESTAV-IPN), 07360 Mexico-City, Mexico.; Max Planck Institute for Molecular Biomedicine, Münster, Germany.; Dept. of Medical Biochemistry, Academic Medical Center, Amsterdam, the Netherlands

**Author notes:** These authors contributed equally (Timo Rademakers, Marieke Goedhart). **Correspondence should be addressed to:** Jaap D. van Buul Molecular Cell Biology Lab, Dept. Plasma Proteins Sanquin Research and Landsteiner Laboratory Academic Medical Center at the University of Amsterdam Plesmanlaan 125 1066CX Amsterdam, the Netherlands.

**Keywords:** Homing, VE-cadherin, Bone marrow, stem and progenitor cell, Permeability, Transmigration

## Abstract

Bone marrow (BM) endothelium plays an important role in homing of hematopoietic stem and progenitor cells (HSPCs) upon transplantation, but surprisingly little is known on how endothelial cells regulate local permeability and HSPC transmigration. We show that temporal loss of VE-cadherin function promotes vascular permeability in BM, even upon low dose irradiation and strongly enhanced homing of transplanted HSPCs to BM of irradiated mice. Intriguingly, stabilizing junctional VE-cadherin *in vivo* reduced BM permeability, but did not prevent HSPC migration into the BM, suggesting that HSPCs enter the bone marrow by transcellularly crossing the endothelium. Indeed, HSPCs induce podosomes to cross human BM endothelial monolayers in a transcellular manner. By contrast, HSPC rather use the paracellular route when VE-cadherin function is inhibited. Taken together, VE-cadherin is crucial for BM vascular homeostasis and HSPC homing, and may therefore serve as a potential therapeutic target to improve HSPC homing strategies.

## INTRODUCTION

Hematopoietic stem cell transplantation (HSCT) is used to restore hematopoiesis in patients with (hematological) malignancies and disorders after chemotherapy and/or irradiation. The first step to therapeutic success of HSCT critically depends on the homing of sufficient numbers of hematopoietic stem-and progenitor cells (HSPC) in the bone marrow (BM) (1). An important step in homing is the actual transmigration of reinfused HPSCs across the BM endothelium to the underlying parenchyma. This extravasation event requires a set of specific molecular interactions that mediate firm adhesion of the HSPCs to the BM endothelial cells, and subsequent transmigration across the endothelial lining (2, 3). Although the role of integrins and other adhesion molecules involved in firm adhesion of HSPCs to the endothelium have been elucidated, and novel *in vivo* imaging approaches are enabling documentation of the dynamic behaviour of HSPC homing to the bone marrow (4), the crucial mechanisms that underlie HSPC transmigration into the BM remain unclear.

The BM is a highly vascularized organ with over 250 blood vessels per square millimeter tissue (5), and these vessels were initially thought to be fenestrated and very permeable (6). However, it has become clear in recent years that there are many hypoxic niches in the BM, especially close to the vasculature (7) (8), which would argue for a tightly regulated vasculature in the BM. Thus, there is still uncertainty about the extent of vascular permeability in the BM under homeostatic conditions.

We have previously shown that blocking VE-cadherin interactions in human BM-derived endothelial cells *in vitro* enhances transmigration of HSPC (9). Moreover, a recent study by Itkin et al. has suggested that extravasation of HSPC correlates with the permeability of different vessel types in the BM (10). Vascular permeability is regulated through the integrity of the endothelial junctions (11): when VE-cadherin contacts are disrupted *in vivo* through administration of anti-VE-cadherin antibodies, vascular permeability in the heart and lung is increased (12). In addition to regulating vascular permeability, VE-cadherin also plays a key role in transmigration of leukocytes (13, 14). VE-cadherin is therefore a likely candidate to control migration of HSPC to the BM. These data suggest that VE-cadherin may function as a target to improve homing conditions of HSPCs after transplantation. Although such an approach would be a highly attractive therapeutic option, it has never been tested under *in vivo* conditions.

We therefore hypothesized that VE-cadherin regulates local BM vascular permeability as well as directional HSPC homing to the BM. Here, we show that *in vivo* blocking of homotypic VE-cadherin interactions increases BM vascular permeability and improves HSPC homing to the BM upon low dose irradiation. Interestingly, we find that HSPCs predominantly cross BM endothelium in a transcellular manner. However, upon loss of VE-cadherin function, HSPCs switch to the paracellular route to cross the BM endothelium. These results show that temporal targeting of VE-cadherin, and thus altering the transendothelial migration mode of HSPCs, may allow more efficient homing of transplanted HSPCs. when VE-cadherin is blocked.

## RESULTS AND DISCUSSION

### Bone marrow vasculature is highly permeable for small molecules

As little is known about the extent of vascular barrier function in the BM during homeostasis, we first set out to determine this. Fluorescently-labeled 10 kDa dextrans were intravenously administered to mice together with the vascular marker GS-I and allowed to circulate for 5 minutes, after which the mice were sacrificed (Fig. 1A). To define vascular permeability, we employed whole mount multiphoton imaging of several organs and measured the fluorescent intensity of the dextrans in the vascular tissue microenvironment within a perimeter of one cell layer (8 μm) around individual blood vessels (Fig. 1B). To correct for potential loss of intensity at greater tissue depth, fluorescence intensity of the dextrans was normalized to intensity values of the GS-I vascular staining (Fig. 1C). We determined the vascular permeability of BM, liver, lung and heart for 10 kDa dextrans as a ratio to the vascular permeability of the brain, where vascular permeability is exceptionally low (Fig. 1D, 1E). Vascular permeability in the BM was comparable to that of the liver, and approximately 2-3 times higher than that of the lung and heart, respectively (Fig. 1D, 1E). Thus, BM vessels are indeed permeable for small dextrans, similarly to the fenestrated sinusoids of the liver. Within the BM, we discriminated between sinusoids and arterioles in the metaphysis and diaphysis area and found, as expected, that basal permeability in sinusoidal venules was significantly higher than in arterioles (Fig. 1F). The niche area where the vessels are located in the BM (meta-and diaphysis) did not affect the permeability scores. These data indicate that baseline permeability differs per type of vessel, but not per niche location in the BM. This high baseline permeability of BM sinusoids may therefore expose adjacent HSPC to blood plasma and increase their reactive oxygen species (ROS) levels, thereby increasing their differentiation and migration capacity (10).

**Figure 1:**
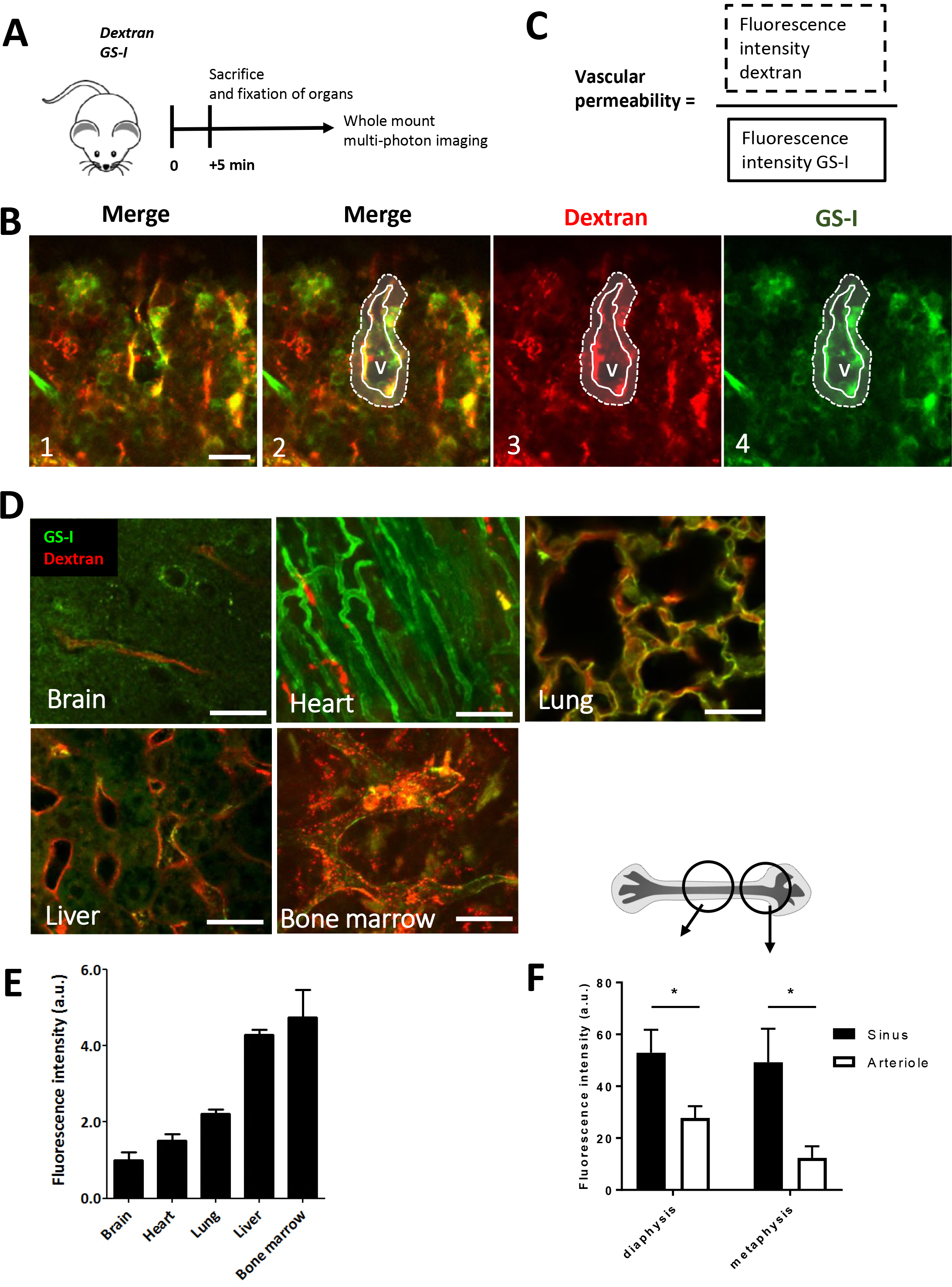
Bone marrow vasculature is highly permeable for small molecules. (**A**) Schematic summary of experimental set up. (**B**) Bone marrow imaging after injection of fluorescent dyes with in green GS-I for vessel labeling and in red 10 kDa Dextran (Panel 1). Panel 2-4 show identification of a vessel (solid line) with red fluorescence detection outside of the vessel (dotted line), as a measurement of vascular permeability. V indicates the vessel lumen. (**C**) Mathematical equation for vascular permeability: Fluorescence intensity of Dextran (solid line in 1B) is divided by the fluorescence intensity of the vessel stain in the selected area (Dotted line in 1B). (**D**) Detection of vascular permeability in several organs as indicated. (**E**) Quantification of the vascular permeability per organ, calculated as described in C (n=2). (**F**) Detailed analysis of vascular permeability in specific bone marrow regions: Diaphysis and metaphysis and discrimination between arterioles and sinusoids, based on intensity of GS-I stainings (n=5). Scale bars indicate 25 μm.

### VE-cadherin regulates BM vascular permeability in homeostatic conditions and after irradiation

As VE-cadherin is recognized as the major regulator of vascular permeability (15), we investigated whether vascular permeability in the BM is regulated by VE-cadherin. To block homotypic binding of VE-cadherin at the endothelial junctions, we injected mice intravenously with a blocking antibody against VE-cadherin (clone 75 (13)) 4 hours prior to the injection of fluorescently labeled 10kD dextran (Fig. 2A). It has previously been shown that VE-cadherin-blocking antibodies increase vascular permeability of heart, lung and lymph nodes *in vivo* (12, 13), but little is known about their effect on the integrity of the BM vasculature. We found that loss of VE-cadherin function resulted in increased vascular permeability in both sinusoids and arterioles in the BM (Fig 2B, 2C) compared to non-treated conditions, indicating that VE-cadherin actively regulates vascular leakage in the BM during homeostasis. Interestingly, when studying the effects of VE-cadherin blockage on the permeability of 500 kDa dextran, no significant increase was detected (Fig 2B, 2C), indicating that the blockage does not massively disrupt endothelial junctions, and showing that VE-cadherin specifically regulates permeability of small molecules through the BM sinusoids in particular.

**Figure 2:**
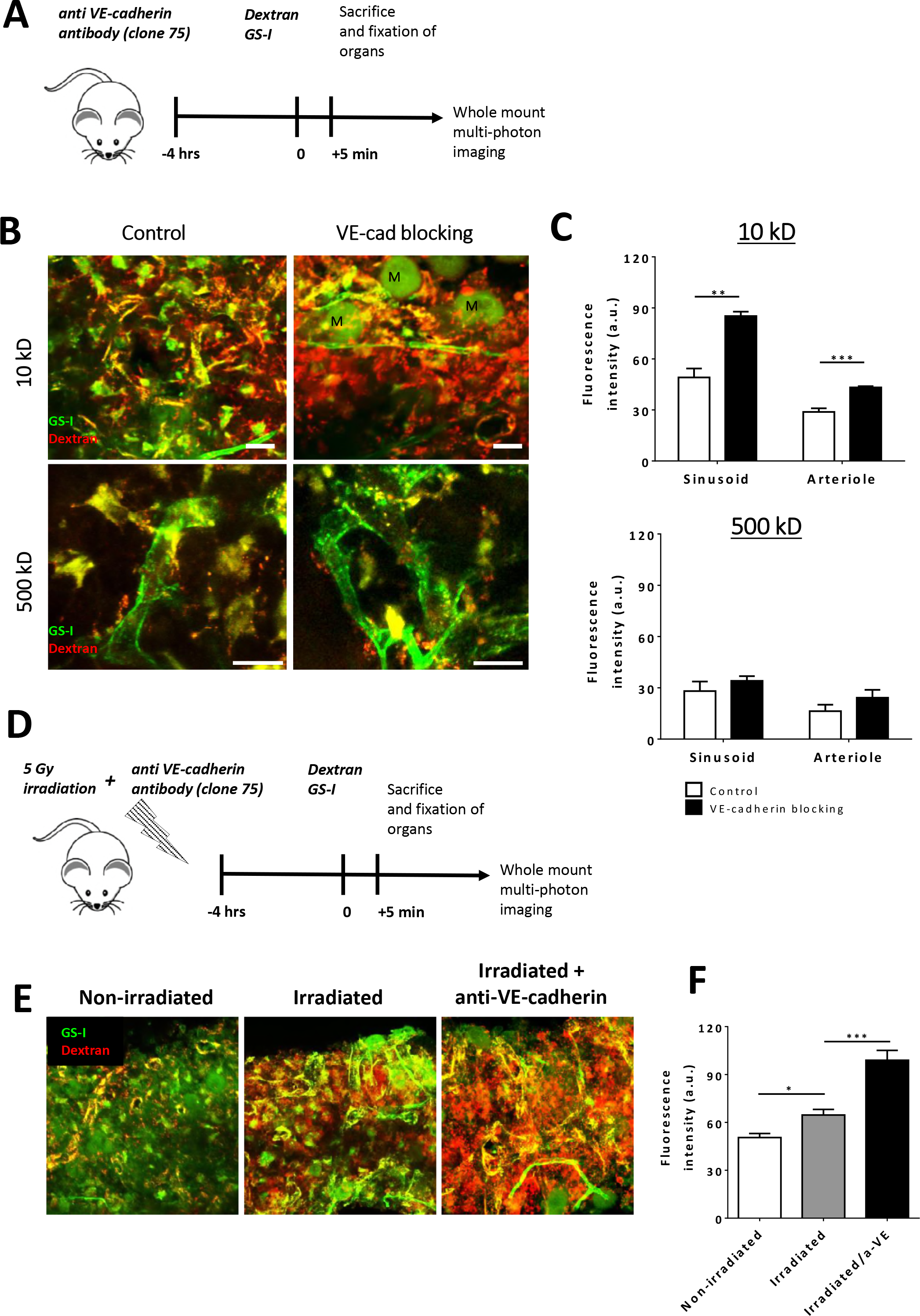
Anti-VE-cadherin antibodies increase BM vascular permeability in homeostatic conditions and after irradiation. (**A**) Schematic summary of experimental set up. (**B**) Bone marrow imaging after injection of fluorescent dyes with in green GS-I for vessel labeling and in red 10 and 500 kDa Dextran as indicated. M: megakaryocytes (**C**) Quantification of vascular permeability in arterioles and sinusoids in metaphysis and diaphysis after blocking VE-cadherin. (10kD: n=7-8 per group, 500kD: n=4 per group) (**D**) Schematic summary of experimental set up including 5 Gy irradiation. (**E**) Bone marrow imaging after treatment as indicated and stained with fluorescent dyes in green GS-I for vessel labeling and in red 10 kDa Dextran. (**F**) Quantification of vascular permeability after irradiation and blocking VE-cadherin (n=4-5 per group). Scale bars indicate 25 μm.

From a clinical perspective, total body irradiation is commonly applied before hematopoietic stem cell transplantation (HSCT) to enable homing of sufficient numbers of HSPC (16). We measured vascular permeability around BM sinusoids of low dose irradiated mice (Fig. 2D) and found that vascular permeability for 10kDa dextrans in the BM is significantly increased after irradiation (Fig 2E, 2F), in line with previous studies (17, 18). Of note, using the VE-cadherin-GFP transgenic knock in mouse model, we did not observe significant changes in vascular morphology in the BM after irradiation, suggesting that the physical structure of the BM vasculature is not affected by irradiation (Fig. S1A). To study in more detail if the molecular complex that underlies the endothelial cell-cell junction integrity is affected upon irradiation, we used endothelial cells isolated from human umbilical cord vein (HUVEC). We found that low dose irradiation induced internalization of VE-cadherin (Fig. S1B, S1C). However, no clear increase in tyrosine phosphorylation of VE-cadherin or a loss of the interaction of p120-catenin with VE-cadherin was observed, although both events are known to be involved in VE-cadherin internalization (19, 20)(Fig. S1D, S1E). From these data we concluded that low dose irradiation does not affect the overall physical structure of endothelial junctions in the BM vasculature, but rather induces a change in the regulation of vascular permeability, possibly by increasing the internalization of VE-cadherin.

Interestingly, although vascular permeability in the BM of mice that were exposed to a low dose irradiation was increased, permeability was even further increased when mice were exposed to a high dose of 10 Gy irradiation (Fig. S2A). This indicates that there is a window to increase BM permeability and potentially homing of HSPC in the default conditioning regimen for HSCT. Therefore, we questioned whether loss of VE-cadherin function additionally enhances vascular permeability, induced by low dose irradiation. Mice were irradiated and injected with VE-cadherin blocking antibody 4 hours prior to administration of 10 kDa fluorescently labeled dextran. As shown in Figure 2E and 2F, loss of VE-cadherin function significantly increased vascular permeability in the BM upon irradiation, demonstrating the functionality of this antibody in the default conditioning regimen for HSCT. Interestingly, we found that while VE-cadherin still controlled BM permeability after low dose irradiation, a high dose of 10 Gy irradiation induces maximal vascular permeability in the BM that is no longer regulated by VE-cadherin (Fig. S2A, S2B).

To substantiate the role of VE-cadherin in regulating the integrity of BM vasculature, we examined permeability in VE-cadherin/α-catenin fusion mice. In these mice, VE-cadherin is genetically replaced by a VE-cadherin-α-catenin fusion construct, resulting in very stable endothelial junctions, because VE-cadherin can no longer dissociate (Fig. 3A) (21). We found that under homeostatic conditions, BM vascular permeability was significantly reduced in VE-α-catenin fusion mice compared to WT controls (Fig. 3B, 3C). The effect of stabilizing endothelial junctions on BM vascular permeability was still apparent after low dose irradiation of VE-α-catenin fusion mice, as BM vascular permeability was significantly lower compared to irradiated controls, in sinusoids as well as in arterioles (Fig. 3B, 3C). Taken together, these data show that VE-cadherin is crucial in regulating vascular permeability in the BM.

**Figure 3:**
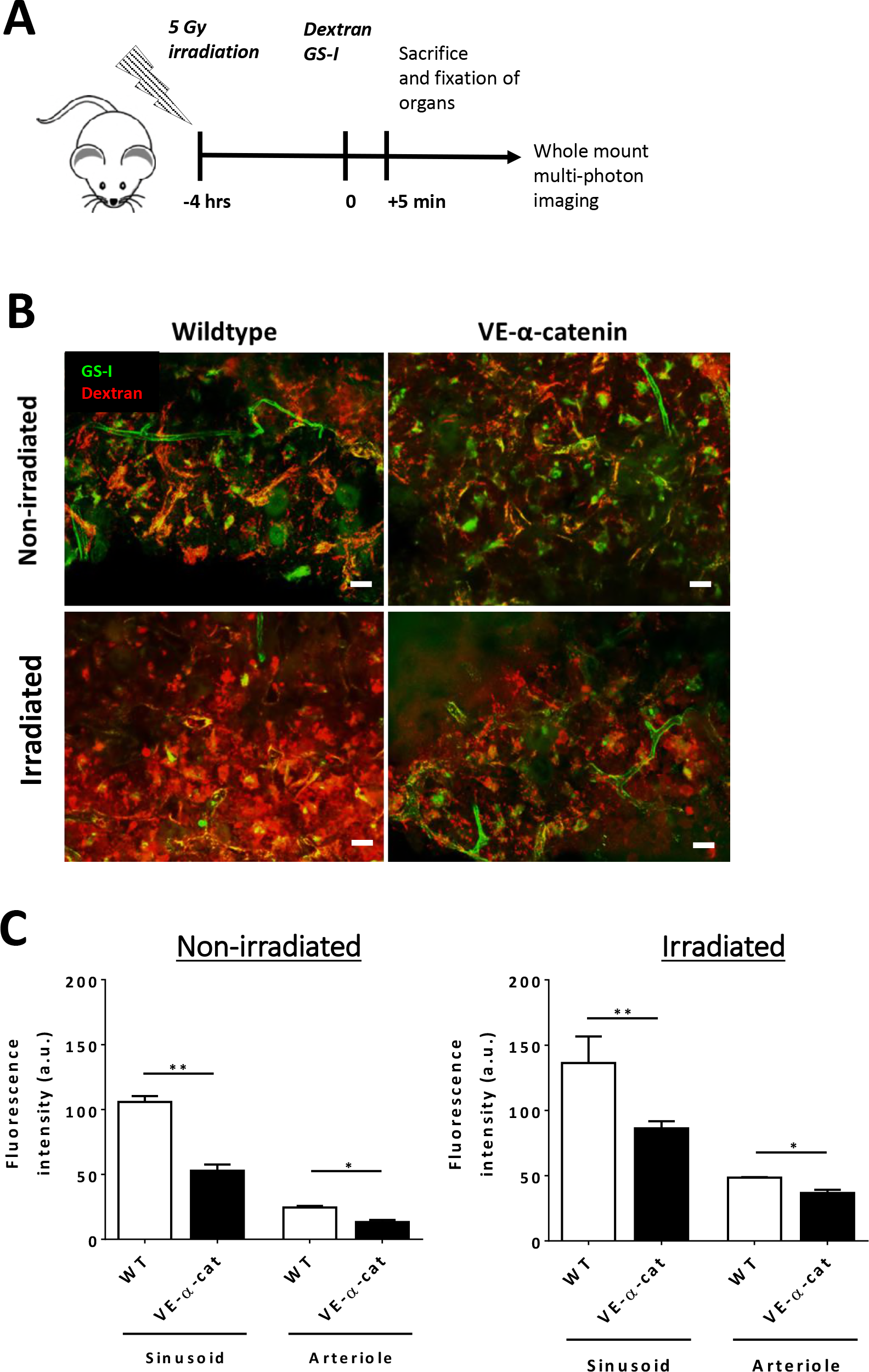
Reduced BM vascular permeability in VE-cadherin-α-catenin fusion mice. (**A**) Schematic summary of experimental set up. (**B**) Bone marrow imaging of wildtype and VE-cadherin-alpha-catenin chimera mice after injection of fluorescent dyes with in green GS-I for vessel labeling and in red 10 kDa Dextran (n=3-4 per group). (**C**) Quantification of vascular permeability in arterioles and sinusoids in wildtype and VE-cadherin-alpha-catenin chimera mice as indicated. Left graph shows results from non-irradiated group and right graph shows results from irradiated group. Scale bars indicate 25 μm.

### Loss of VE-cadherin function increases HSPC homing to the BM

We next examined whether VE-cadherin also regulates homing of HSPC to the BM. Lineage^−^Sca-1^+^c-kit^+^ (LSK) cells were adoptively transferred into low dose-irradiated mice in the presence or absence of a blocking VE-cadherin antibody (Fig. 4A). After 16 hours, mice were sacrificed and the presence of donor HSPC in BM, lung, spleen, and liver was determined (Fig. 4B). We found that homing of LSK cells to the BM was significantly increased 2-fold in mice treated with anti-VE-cadherin antibody compared to controls (Fig. 4C). Of note, there was also a tendency towards increased homing of donor HSPC to spleen in anti-VE-cadherin injected mice albeit not significant (Fig. 4C). Homing of HSPC to lung and liver was not affected by anti-VE-cadherin antibodies. The reason for this may be that percentages of donor HSPC in other organs than the BM were >40 fold lower than in the BM (data not shown), implying that the presence of adoptively transferred HSC in these organs was not due to directed migration. This is not surprising considering that CXCL12, the most important chemokine for HSPCs, is expressed in the BM at much higher levels than in any other organ (22). Thus, in addition to regulating vascular permeability in the BM, VE-cadherin regulates the directed migration of HSPC to the BM.

**Figure 4:**
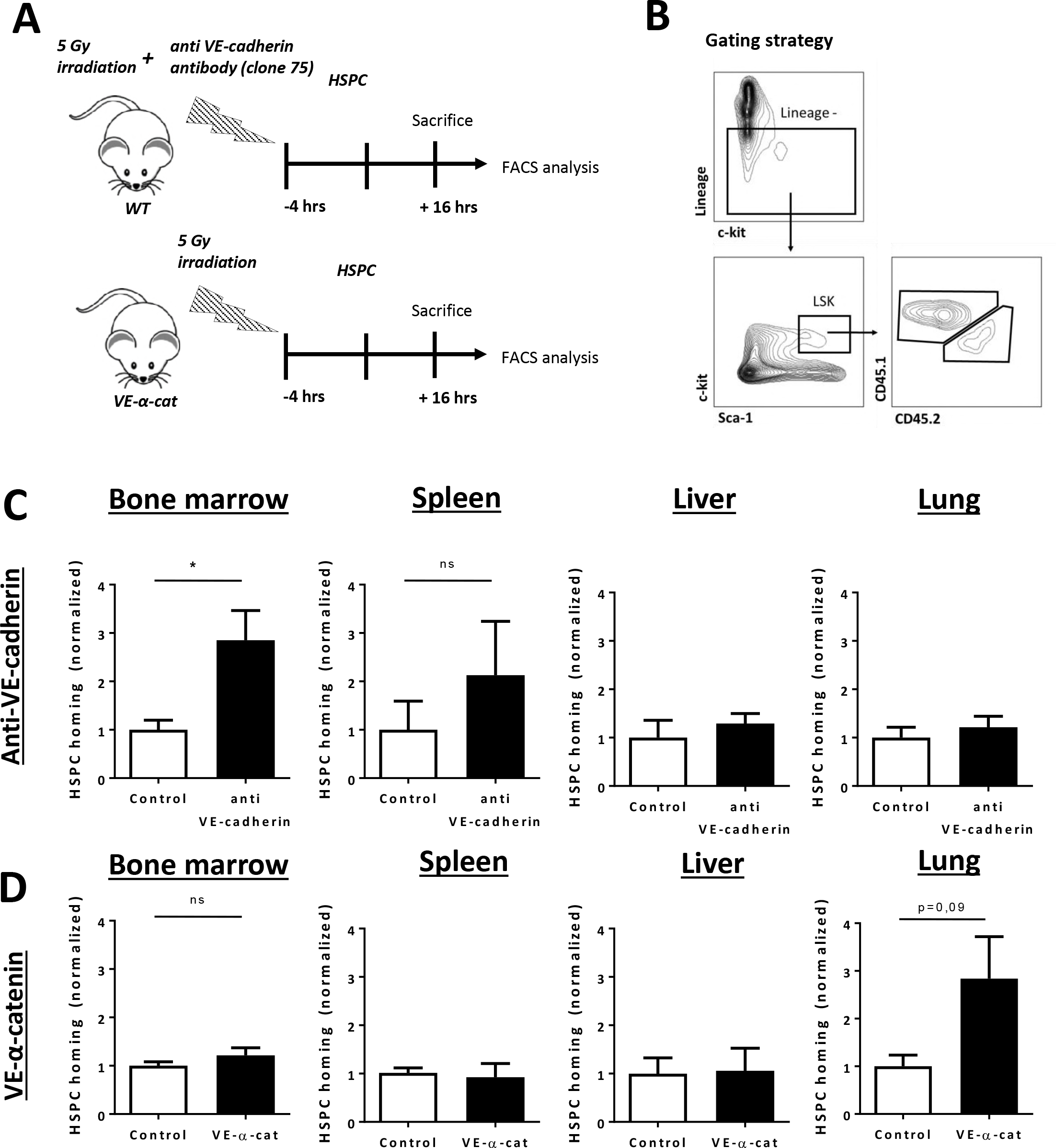
Anti-VE-cadherin antibodies increase HSPC homing to the BM. (**A**) Schematic summary of experimental set up for three groups: wildtype (WT), anti-VE-cadherin antibody administration and VE-cadherin/alpha-catenin chimera (VE-α-cat). (**B**) Gating strategy to select for LSK cells (murine HSPCs) using flow cytometry. (**C**) Normalized homing efficiency of HSPCs to different organs after 16 hours of transplantation in control and anti-VE-cadherin antibody group (n= 4-5 per group, representative of two independent experiments). (**D**) Normalized homing efficiency of HSPCs to different organs after 16 hours of transplantation in WT and VE-α-cat groups (n= 4-5 per group).

As we observed a clear correlation between increased BM vascular permeability and increased homing of HSPC to the BM upon loss of VE-cadherin function, we expected decreased homing of HSPC to the BM of VE-cadherin-α-catenin fusion mice that have decreased BM vascular permeability. Surprisingly, we did not observe decreased homing of HSPC to the BM of VE-cadherin-α-catenin fusion mice compared to WT controls (Fig 4D), indicating that permeability and transmigration are regulated separately. This is in line with our previous finding where we describe a specific role for endothelial cortactin in both permeability and neutrophil recruitment (23). There was no difference in HSPC homing to lung, spleen, and liver in VE-α-catenin fusion mice (Fig. 4D). To verify that the donor HSPC that we identified in the BM of VE-α-catenin fusion mice by FACS analysis were not trapped in the vascular bed of the BM, cryosections of the BM of transplanted wildtype and VE-cadherin-α-catenin fusion mice were analyzed (Fig. S2C). Here, we found that the majority of adoptively transferred HSPC were located inside the BM parenchyma (Fig. S2C, S2D), confirming that stabilizing endothelial junctions does not impair transendothelial migration of HSPC into the BM.

### HSPC predominantly use the transcellular route of migration over BM endothelium and form podosome-like structures

Our surprising findings of increased HSPC homing upon loss of VE-cadherin function, but unchanged homing in VE-cadherin-α-catenin fusion mice, led us to examine the transendothelial migration route that HSPCs use to enter the BM in more detail. Leukocytes can traverse the endothelial barrier by two distinct routes: paracellularly, i.e. through VE-cadherin-based cell-cell junctions and transcellularly, i.e. directly through individual endothelial cells, leaving the endothelial junctions intact (24, 25). To determine whether HSPCs predominantly take the para-or transcellular route under homeostatic conditions, we employed an *in vitro* migration assay using human CD34^+^ cord blood (CB) HSPCs and immortalized human bone marrow endothelial cells (HBMEC) (26). To properly discriminate between the two routes, HBMECs were transfected with Lifeact-GFP to visualize F-actin, activated for 4 hours with IL-1β (26) and 30 minutes prior to adding the CD34+ cells, HBMECs were incubated with fluorescently-labeled, non-blocking VE-cadherin antibodies to visualize endothelial junctions (Fig. 5A) (27). We found both para-and transcellular migration events of HSPC that crossed HBMECs. Quantification showed that the vast majority (~75%) of HSPC cross the HBMEC transcellularly (Fig. 5B). Interestingly, administration of a blocking VE-cadherin antibody prior to the addition of the HSPCs to the HBMECs resulted in a significant increase of HSPCs that migrated paracellularly (Fig. 5B). To show the importance of transcellular migration events in HSPC migration across BM endothelium, we blocked membrane fusion activity in endothelial cells, known to prevent the formation of transcellular pores (28). HSPC migration across a confluent HBMEC layer that was treated with the SNARE-complex inhibitor N-Ethylmaleimide (NEM) showed a significant inhibition of the number of HSPCs transmigrating towards CXCL12 (Fig. 5C), indicating that the formation of transcellular pores is the major route for HSPCs to cross HBMEC. This is a surprising finding considering that leukocytes mostly use the paracellular route for transmigration (29). Although the exact mechanisms of diapedesis are still under debate, trans-and paracellular migration are most likely regulated by different molecular mechanisms. On the endothelial side, VE-cadherin plays a key role in paracellular migration (30), On the other hand, Mac-1/CD11b on neutrophils is thought to be essential for crawling towards the nearest endothelial junction, thus facilitating paracellular migration (31). Interestingly, only a small subset of HSPCs express Mac-1/CD11b (32), suggesting that the majority of HSPCs is incapable to efficiently locate the endothelial junctions. To test this hypothesis, we used *in vitro* physiological flow conditions and live cell imaging (27) to quantify crawling of neutrophils and HSPCs over HBMEC monolayers. Neutrophils crawled extensively on HBMEC monolayers, as they did on HUVECs (27)(27)(Kroon et al., 2014)(Kroon et al., 2014)(Kroon et al., 2014)(Fig. 5D; Fig. S3A-C). In contrast to neutrophils, HSPCs did not crawl on HBMEC monolayers under physiological flow conditions, but rather remained sessile upon adhesion (Fig. 5D-F). Thus, our data suggest that HPSC are not equipped to locate the nearest endothelial junction upon their attachment, and therefore require the transcellular pathway for endothelial transmigration.

**Figure 5:**
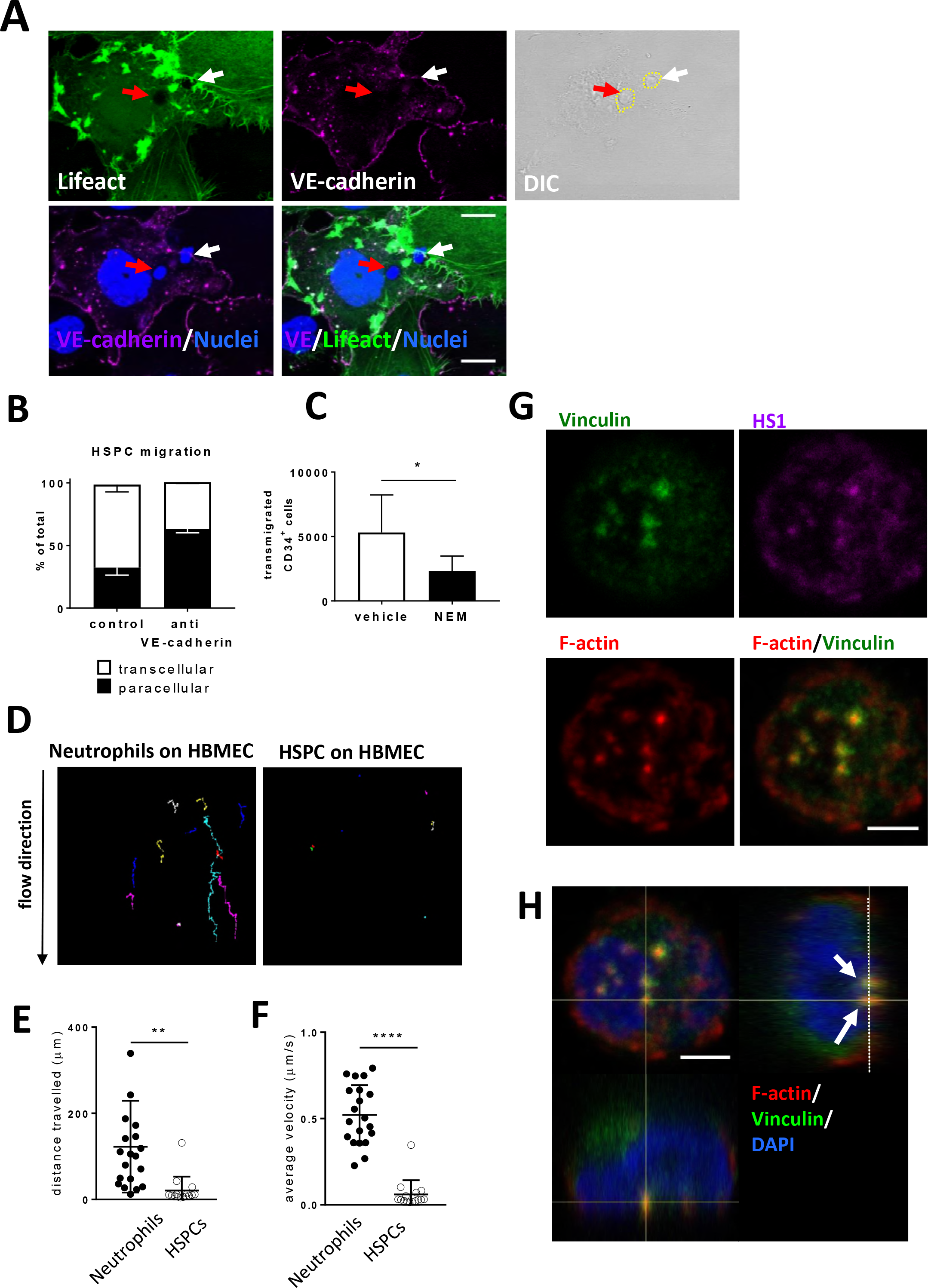
HSPC predominantly use the transcellular route of migration across BM endothelium by forming podosome-like structures. (**A**) Human bone marrow endothelial cells (HBMEC) were cultured and pretreated with IL1β and transfected with GFP-LifeAct to monitor actin dynamics during HSPC interactions. Both migration routes were detected for HSPC migration over HBMEC: paracellular, i.e. through the cell-cell junctions, illustrated by the white arrow, and transcellular, i.e. through the endothelial cell body, illustrated by the red arrow. VE-cadherin in magenta marks endothelial cell-cell junctions, Nuclei are stained with Hoechst in blue. Yellow dotted circles indicate localization of the human HSPCs. Scale bar indicates 20μm. (**B**) Quantification of transmigration route of HSPCs. Open bar represents percentage of cells that cross transcellularly and closed bars represent percentage that crosses paracellularly (Mean ± SD, n=3 per group). (**C**) Blocking of membrane fusion events in HBMEC by N-Ethylmaleimide (NEM) impairs HSPC transendothelial migration towards CXCL12 in an *in vitro* Transwell system (Mean ± SD, n=3 per group). (**D**) Migration tracks of cells under flow conditions show highly motile neutrophils and passive HSPCs, quantified by distance travelled on the apical surface of HBMEC in μm (**E**) or average velocity (μm/s) **(F).** These experiment were independently carried out at least three times. (**G**) HSPCs were treated for 30 minutes with PMA showing the induction of podosomes, based on typical podosome markers: vinculin (green) in the circle and HS1 (magenta) and F-actin (red) in the core. Scale bar indicates 3 μm (**H**) Orthogonal projection shows the distribution of the podosomes at the basolateral surface (arrows) with F-actin in red, vinculin in green and the nucleus in blue.

As of yet, little is known about the site of transmigration of HSPC (33). A prerequisite for transcellular migration of leukocytes is the formation of invasive protrusions, or ‘podosomes’, to palpate the surface of the endothelium and initiate transcellular pore formation (28). Podosomes consist of a protrusive actin core and an adhesive ring of integrins and adaptor proteins, such as vinculin and cortactin, or its hematopoietic homologue, hematopoietic lineage cell-specific protein 1 (HS-1) (34, 35). To determine whether HSPCs can form podosomes, we allowed HSPCs to adhere on fibronectin-coated coverslips and exposed them to PMA for 30 minutes. As a positive control we induced podosome formation in cultured dendritic cells, where we indeed observed clusters of podosomes with a ring of vinculin and a core containing actin and HS-1, as previously described ((35); Fig. S3D, S3E). Remarkably, we could identify similar, podosome-like structures in HPSCs (Fig. 5G, 5H), indicating that HSPCs are equipped with the machinery to induce podosomes to migrate through endothelial cells in a transcellular manner.

We provide here for the first time evidence that HSPCs can form podosome-like structures and that diapedesis of HSPCs is predominantly transcellular under homeostatic conditions. In addition, a higher proportion of HSPCs use the paracellular route when junctional integrity is diminished upon VE-cadherin blocking, indicating that junctional resistance also plays a role in the molecular decision for HSPC transmigration. Putatively, transcellular migration of HSPCs remains upon administration of a VE-cadherin antibody, and the observed increase in HSPC homing to the BM can be explained by an increase of paracellular migration of HSPCs. The fact that HSPCs predominantly use the transcellular route of transmigration can also explain that HPSC homing to the BM is not impaired in VE-cadherin-α-catenin fusion mice. It was previously shown that, in contrast to our data on HSPC homing, neutrophil migration to inflamed cremaster, lung, and skin is significantly impaired in VE-cadherin-α-catenin fusion mice (21). However, this is not surprising considering that neutrophils almost exclusively use the paracellular route of transmigration (29, 31, 36, 37).

Although we have shown that anti-VE-cadherin antibodies increase trafficking of HSPCs to the BM, a potential side effect of disturbing the integrity of the endothelial barrier in the BM is an elevation of reactive oxygen species (ROS) levels in the BM. Itkin and colleagues have recently shown that disruption of the endothelial barrier using genetically engineered mice, similarly to anti-VE-cadherin antibodies, increases bi-directional trafficking of HSPC to and from the BM (10). They also showed that increased ROS levels upon continuous disruption of the endothelial barrier hampered maintenance of HSPCs. However, proper timing of administration of anti-VE-cadherin antibodies before HSCT likely limits potential side effects, as the integrity of the endothelial barrier is most likely restored once HSPCs have homed to the BM. In favor of this view, we show that BM vascular permeability is back to baseline levels 16 hours after *in vivo* administration of anti-VE-cadherin antibodies (Fig. S2E).

There are several potential clinical benefits from using anti-VE-cadherin antibodies to increase HSPC homing to the BM upon HSCT. Increasing the proportion of HSPCs that home to the BM most likely advances immune reconstitution, which is pivotal for survival of HSCT patients (38). This is especially important for small hematopoietic grafts such as cord blood derived HSPCs, which have a high suitability for recipients and are readily available, but are currently the last choice of hematopoietic graft due to the limited amount of CD34^+^ cells that can be harvested from one umbilical cord (39). Similarly, anti-VE-cadherin antibodies may aid homing of *ex vivo* expanded and (genetically) manipulated HSPCs, which often have reduced repopulating potential (40). Furthermore, administration of anti-VE-cadherin antibodies may lower the radiation dose that is needed for adequate homing of HSPCs upon HSCT, and as such reduce irradiation side-effects. Importantly, injection of anti-VE-cadherin was well tolerated and did not lead to any overt side-effects in other organs. However, increasing the concentration of VE-cadherin blocking antibodies likely leads to morbidity and/or mortality, as has been shown previously (12).

Taken together, we report that although HSPCs predominantly use the transcellular route for transendothelial migration in homeostasis, blocking VE-cadherin homotypic interactions favors paracellular migration of HSPCs and increases homing of HSPCs to the BM. Our work offers valuable insight into the mechanisms of HSPC homing to the BM, and we have identified VE-cadherin as a potential target to increase the success of HSCT.

## METHODS

### Mice

The following strains were used: C57BL/6, C57BL/6-Ly5.1, VE-cadherin/α-catenin (21) and VE-cadherin-GFP (41). Mice were maintained on a C57BL/6 background in the animal facilities of the Netherlands Cancer Institute (Amsterdam, The Netherlands) and the Max Planck Institute for Molecular Biomedicine (Münster, Germany) in specific pathogen-free conditions. All animal experiments were approved by the local Animal Ethical Committee in accordance with national regulations.

### Multi-photon imaging of vascular permeability

Mice were injected with 200 μl PBS containing the vascular marker GS-I (42) (final concentration 30 μg/ml; Vector Labs), and fluorescently labelled fixable dextran 10kD and/or 500kD (final concentration 30μg/ml; Thermo Fisher Scientific) This was allowed to circulate for 5 minutes before the mice were euthanized, after which tissue (BM, heart, lung, brain, liver, spleen) were dissected and fixed for 4 hours in 4% paraformaldehyde. The tissues were kept in PBS at 4°C before whole mount imaging using multiphoton microscopy. For measuring permeability upon VE-cadherin blocking, similar procedure was followed, yet four hours prior to injection with dextrans and GS-I, mice were injected with a VE-cadherin blocking antibody (13) (6 μg/mouse; clone 75; BD Transduction Laboratories). For measuring permeability upon irradiation, mice were irradiated (4 or 10 Gy, depending on the experiment) 4 hours prior to dextran injection. Where indicated, mice were also injected with VE-cadherin blocking antibody directly after irradiation as described above. Multiphoton imaging was performed on a Leica SP8 system equipped with a Spectraphysics Insight Deepsee laser and a 25× 0.95NA water immersion objective (Leica Fluotar VISIR). Excitation wavelength was tuned to 880nm, and NDD HyD detectors were used for detecting emission for FITC (525/50nm) and Texas Red (585/40nm). For 3-colors sequential scanning was performed detecting far red (AlexaFluor633) at 650-700nm using a regular PMT. Imaging was performed at 12-bit, at a frame size of 1024×1024 pixels (pixel size: 0.435 × 0.435μm), scanning at 300Hz and an interplanar distance of 0.50-1.00μm. Microvascular permeability was assessed using ImageJ (http://imagej.nih.gov/ij/), and was determined by calculating the fluorescence intensity of dextrans outside the microvessel (extraluminal area) in the various tissues. Extraluminal area was defined using a one cell layer cutoff (approximately 6um), as was assessed experimentally by measuring diffusion distance at baseline by performing line scanning (not shown). To correct for loss of intensity at greater tissue depth, fluorescence intensity of the dextrans was normalized to intensity value of GS-I vascular staining. All values were background-corrected. The mean ratio was calculated per time point and plotted.

### Murine HSPC homing assay

Murine BM cells were obtained by crushing femurs and tibiae with a mortar and pestle, and the cell suspension was filtered through a 40-μm cell strainer. Murine c-kit+ HSPC were isolated with c-kit microbeads (StemCell Technologies). HSPC were thoroughly washed in PBS and injected intravenously into recipient mice irradiated with a dose of 5 Gy. 16h after HSPC transfer, recipient mice were euthanized and homing of HSPC to BM, spleen, lung, and liver was determined with flow cytometric analysis.

### Flow cytometry

Murine BM cells were obtained by crushing femurs and tibiae with a mortar and pestle. Single splenocyte suspensions were prepared by crushing the spleen through a 70-μm cell strainer with the plunger of a syringe. For flow cytometric analysis of spleen and BM T cells, single cell suspensions were stained for lineage markers (CD4 (clone GK1.5), CD8 (clone 53-6.7), B220 (clone RA3-6B2), CD11b (clone M1/70), Gr-1 (clone RB6-8C5) and Ter119 (clone TER-119)), Sca-1 (clone D7), c-kit (clone 2B8), CD45.1 (clone A20) and CD45.2 (clone104) for 30 minutes at 4°C. Antibodies were obtained from eBioscience, unless indicated otherwise. Flow cytometric analysis was perfomed on a FACSCanto II (BD Biosciences).

### Confocal microscopy of murine BM sections

Bones from VE-cadherin/α-catenin mice were mounted directly in tissue tek embedding compound (Sakura Finetek) and snap frozen in liquid nitrogen. Bones from VE-cadherin-GFP mice were treated for 4 hours with 2% PFA prior to mounting in tissue tek and snap freezing. 8μm cryosections were prepared using the CryoJane^®^ Tape-Transfer System (Leica). Sections were air-dried, sections from unfixed bones were fixed for 10 minutes in dehydrated acetone and all sections were blocked with 5% BSA/PBS. Antibodies used for immunolabelling of BM sections were CD45.1 (clone A20; Biolegend) and CD144 (VE-cadherin; clone BV13; Biolegend). In some cases, sections were counterstained with DRAQ5 (ThermoFisher Scientific). Sections were mounted with Mowiol and imaged using a confocal microscope (Leica TCS SP8)

### Human HSPC migration assays

Cord blood (CB) was collected according to the guidelines of Eurocord Nederland and CD34^+^ cells were isolated as previously described (43). Generation of HBMEC cell lines was previously described (26), and HBMEC were cultured in complete EGM-2 MV medium (Lonza). For static migration assays, HBMEC were cultured until confluency on coverslips or 5 μm transwell insert filters, stimulated for 16h with 10 ng/ml IL-1β (Preprotech), and washed 2 times with assay medium (IMDM (Lonza) containing 10% FCS). In transwell migration assays, 100 ng/ml recombinant human CXCL12 (Strathmann Biotech) was added in assay medium in the lower compartment. In some conditions, HBMEC were pre-treated with 10μg/ml α-VE-cadherin antibody (clone 75; BD Transduction Laboratories) for 30 minutes, and α-VE-cadherin antibody remained present during the transendothelial migration assay. In some conditions, HBMEC were pre-treated with NEM (300 μM; Sigma) or equal dilutions of vehicle (EtOH) for 1 minute, and washed 2 times with assay medium. Subsequently, HBMECS were co-cultured with CD34^+^ HSPC for 3 hours. In static migration assays, cells were fixed using 4% PFA for 10 minutes, washed in PBS and coverslips were mounted with Vectashield with DAPI (Vector Labs) on microscope slides. Para-or transcellular migration of HSPC was determined using a confocal microscope (Leica TCS SP8). In transwell migration assays, migration of HSPC to the lower compartment was quantified by flow cytometry using Cyto-Cal Count Control beads (ThermoFisher Scientific).

### Physiological flow assays

Physiological flow experiments were performed as previously described (27). Briefly, HBMEC and HUVEC were cultured in a FN-coated ibidi m-slide VI0.4 (ibidi, Munich, Germany), HBMEC were stimulated for 16h with 10 ng/ml IL-1β (Peprotech) and HUVEC were stimulated for 16 hours with 10 ng/ml TNFα (Peprotech). CD34^+^ HSPC and freshly isolated neutrophils were resuspended in HEPES medium (20mM HEPES, 132mM NaCl, 6mM KCL, 1mM CaCL2,1mM MgSO4, 1.2mM K2HPO4, 5mM glucose (all from Sigma-Aldrich), and 0.4 % (w/v) human serum albumin (Sanquin Reagents), pH7.4) and were incubated for 30 min at 37 °C. Ibidi flow chambers were connected to a perfusion system and exposed to 0.2 - 0.5 ml/min HEPES shear flow. CD34^+^ HSPC or neutrophils were subsequently injected into the perfusion system and real-time leukocyte-endothelial interactions were recorded for 20 min by a Zeiss Observer Z1 microscope using a 40x numerical aperture (NA) 1.3 oil immersion objective. Velocity of crawling leukocytes and distance travelled over the endothelium were manually quantified using the ImageJ plug-in Cell Tracker.

### Podosome formation assays

For immunofluorescence of podosome formation, CD34^+^ HSPC were grown on fibronectin-coated 12-mm coverslips. As a control, human DCs were generated from peripheral blood mononuclear cells as described previously (44) and also grown on fibronectin-coated 12-mm coverslips. Cells were stimulated with phorbol 12-myristate 13-acetate (PMA; Sigma) for 30 minutes to induce podosome formation. After treatment, the cells were washed in cold PBSA (PBS, 0.5% (w/v) BSA, 1 mM CaCl2, 0.5 mM MgCl2) and fixed in 4% (v/v) formaldehyde in PBS (supplemented with 1 mM CaCl2, 0.5 mM MgCl2) for 10 min. After fixation, the cells were permeabilized in PBS-T (PBS + 1% [v/v] Triton X-100 and 10% [v/v] glycerol) for 4 min, followed by a blocking step in PBS supplemented with 2% (w/v) BSA. The cells were incubated with primary and secondary antibodies as indicated, and after each step washed 3 times in PBS-A. Coverslips were mounted with Vectashield with DAPI (Vector Labs) on microscope slides and imaged with a confocal microscope (Leica TCS SP8).

### VE-cadherin internalization assays

HUVEC were cultured on fibronectin coated glass coverslips, and subjected to 4 Gy irradiation or left untreated. 30 minutes prior to irradiation, a FITC-labelled VE-cadherin antibody (polyclonal; eBioscience) was added to the medium. Cells were fixed at 10, 30, and 60 minutes after irradiation using 4% PFA for 10 minutes, washed in PBS, and stained with an AlexaFluor 647 labelled VE-cadherin antibody (clone 55-7H; BD BD Biosciences) without permeabilizing to stain the membrane fraction. Coverslips were mounted with Vectashield with DAPI (Vector Labs) on microscope slides and imaged with a confocal microscope (Leica TCS SP8). Images were analyzed using ImageJ by determining the colocalization coefficient of VE-cadherin on the membrane, and by determining the fluorescence intensity of VE-cadherin intracellularly.

### Immunoprecipitation and western blot analysis

Immunoprecipitation and western blot were performed as previously described (45). Briefly, HUVEC were cultured in a 10cm2 dish and subjected to 4 Gy irradiation for 1 h or left untreated. Cells were lysed in cold NP-40 lysis buffer, and incubated with 0.5 μg of mAb to VE-cadherin (clone BV6, Millipore) and 50 μl of Dynabeads (Invitrogen) at 4°C with continuous mixing. Subsequently, beads were washed with NP-40 lysis buffer and boiled in SDS sample buffer containing 4% β-mercaptoethanol. Samples were analyzed by SDS-PAGE. Proteins were transferred onto a 0.2 μm nitrocellulose membrane (Whatman, Dassel, Germany), and incubated with specific primary antibodies against p120-catenin, beta-catenin, actin and phosphotyrosine (pY-20) (Abs from Transduction Labs) overnight at 4°C, followed by incubation with secondary HRP-labeled antibodies for 1 h at room temperature. Staining was visualized with an enhanced chemiluminescence (ECL) detection system (ThermoScientific, Amsterdam, The Netherlands). Alternatively, blots were incubated with IR-680-or IR-800-dye-conjugated secondary antibodies. The infrared signal was detected and analyzed using the Odyssey infrared detection system (Li-cor Westburg).

### Statistics

Mean values plus or minus standard error of the mean (SEM) are shown, unless indicated otherwise. For group comparisons, data were tested for Gaussian distribution, after which a Student t-test (Gaussian) or Mann-Whitney U test (non-Gaussian) was used to compare individual groups; multiple groups were compared by ANOVA or Kruskall-Wallis tests, with Bonferroni or Dunn’s post-hoc test, respectively. Statistics were performed using Graphpad Prism 5.0. *P < 0.05; **P < 0.01; ***P < 0.001.

## ACKNOWLEDGEMENTS

The authors thank Simon Tol, Erik Mul, and Aafke de Ligt for technical assistance and the staff of the animal facilities of the NKI and the Max Planck Institute for Molecular Biomedicine for excellent animal care. The authors thank Alessandra Cambi and Koen van den Dries for sharing their expertise on podosomes. AGP received a pre-doctoral scholarship from the Mexican National Council for Science and Technology (Conacyt, 369767); and a travel fellowship grant from the Journal of Cell Science, The Company of Biologists Limited (www.biologists.com) to realize a research stay in the lab of JDvB. Work in the lab of MS is funded by Conacyt (grant 233395). This work was supported by Sanquin Research (PPOC13-030P grant). The authors declare that they have no conflict of interest.

## Legends Supplemental figures

**Supplemental figure 1:**
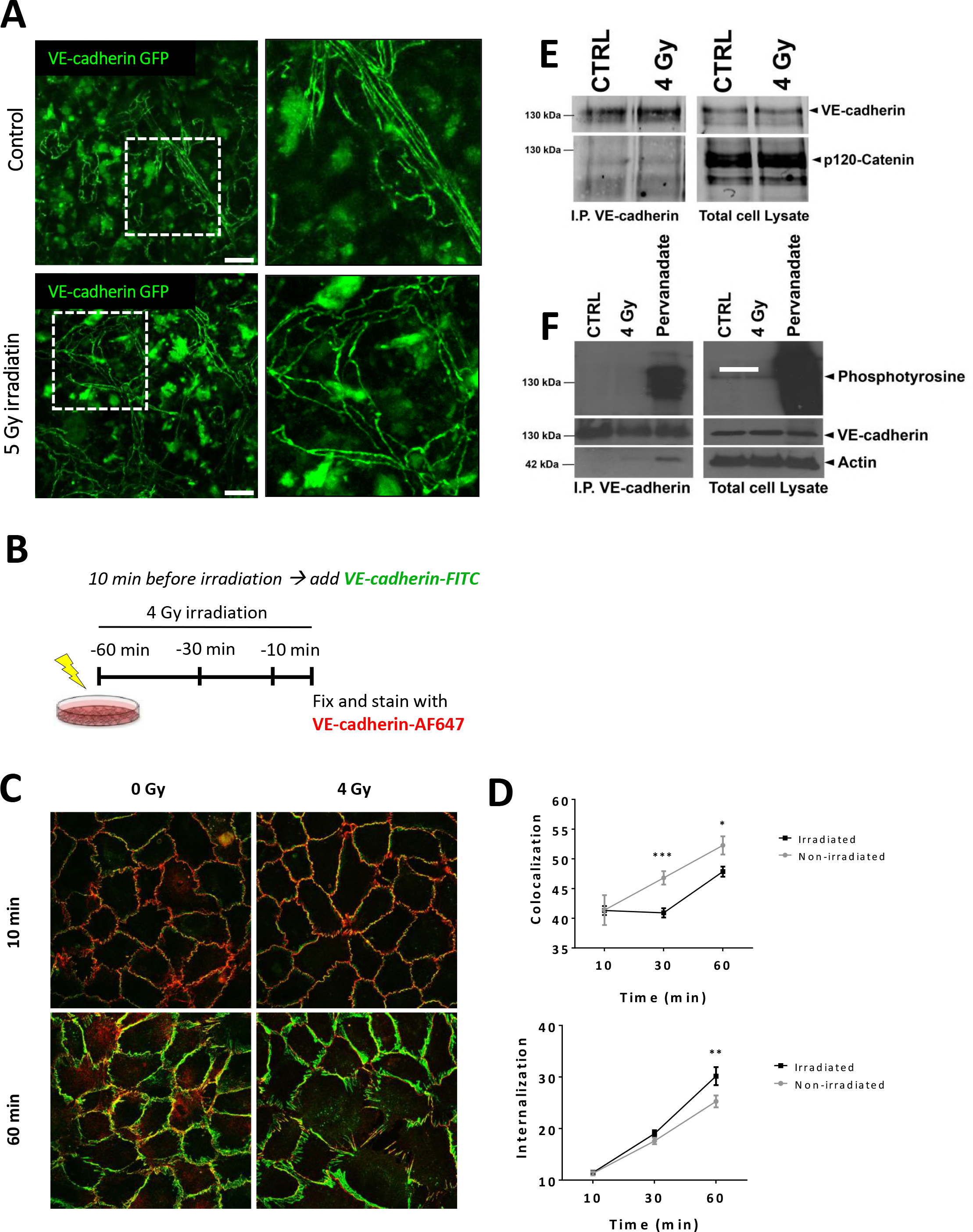
The effect of irradiation on the VE-cadherin complex. (A) Bone marrow imaging of VE-cadherin-GFP transgenic knock in mice after low dose total body irradiation (TBI) or left untreated (Control). Zoom shows intact vessels after sTBI. (n=4 per group, representative image is shown) (B) Schematic summary of experimental set-up for C and D (C) HUVECs were cultured on fibronectin coated glass covers and pre-incubated with a VE-cadherin antibody (red) prior to 4 Gy treatment or left untreated, as indicated; followed by additional VE-cadherin staining (green) on fixed but not permeabilized cells, making it possible to discriminate between internalized VE-cadherin and membrane VE-cadherin. (D) Quantification of membrane-bound VE-cadherin, showing that 4 Gy induces loss of membrane localization of VE-cadherin, measured by co-localization of both VE-cadherin antibodies, as explained in B. Lower graph shows quantification of internalized VE-cadherin, showing that 4 Gy induces VE-cadherin internalization after 60 minutes. Experiment was independently carried out at least three times. (E) VE-cadherin was precipitated from HUVEC cell lysates after 60 minutes of 4 Gy treatment or left untreated and analyzed for p120-catenin binding. Although similar amounts of VE-cadherin are precipitated in both conditions, no difference for p120-catenin binding is detected. (F) VE-cadherin was precipitated from HUVEC cell lysates after 60 minutes of 4 Gy treatment or left untreated and analyzed for changes in tyrosine phosphorylation levels. As positive control, HUVECs were treated for 5 minutes with pervanadate. Although similar amounts of VE-cadherin are precipitated in all conditions, no difference for tyrosine phosphorylation of VE-cadherin is detected. Pervanadate massively increases the phosphotyrosine levels of VE-cadherin (left panel) and other proteins (right panel; total cell lysates). Actin is shown as protein control in the total cell lysates. Experiment was independently carried out at least three times.

**Supplemental figure 2:**
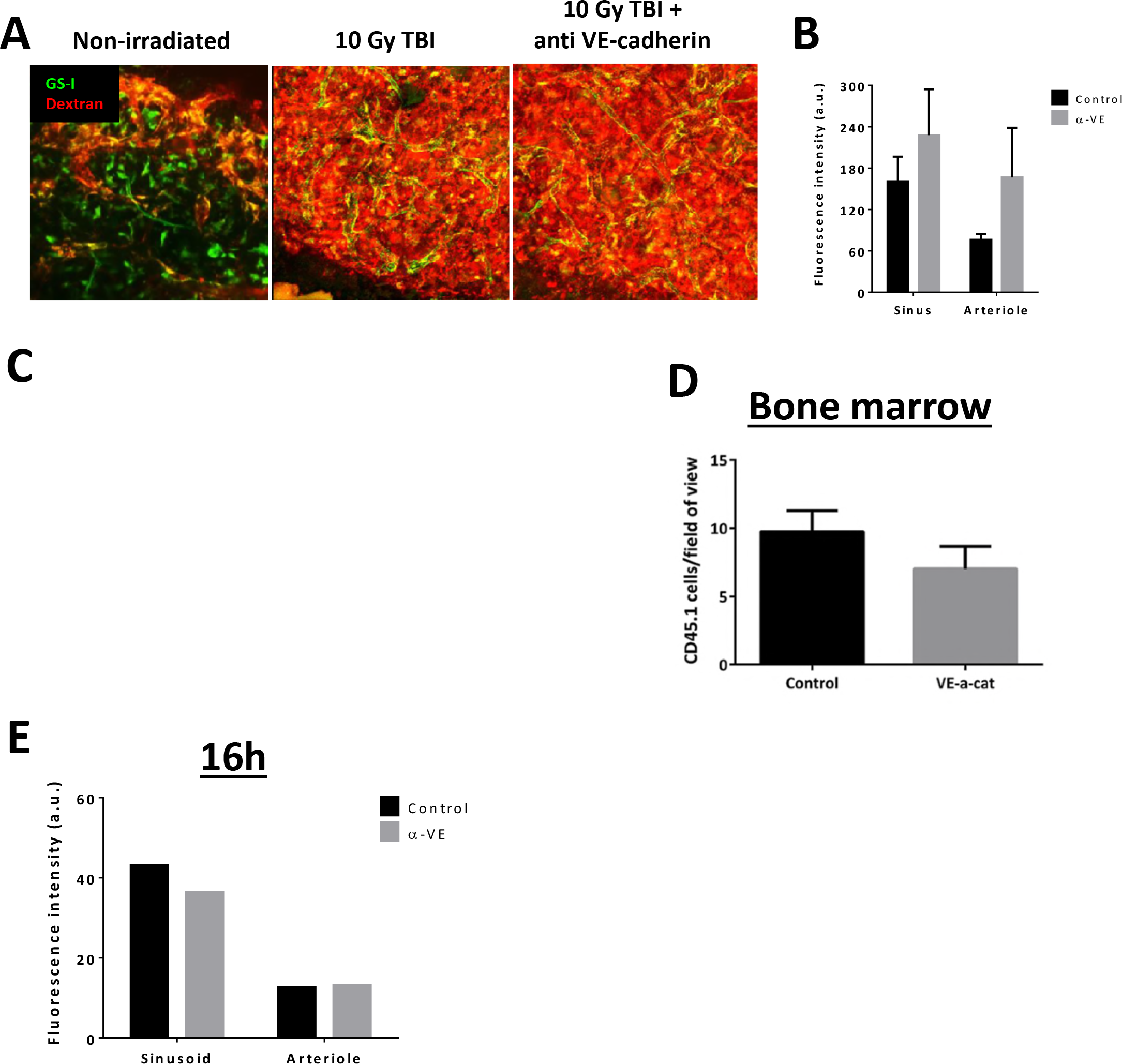
The role of VE-cadherin in BM permeability and homing of HSPC after total body irradiation. (A) Bone marrow imaging after treatment with 10 Gy irradiation and blocking VE-cadherin antibody as indicated, injection of fluorescent dyes with in green GS-I for vessel labeling and in red 10 kD Dextran. (B) Quantification shows no significant increase in vascular permeability after 10 Gy total body irradiation (TBI) when VE-cadherin is blocked (n=5 per group). (C) Imaging of the bone marrow sections from HSPC homing experiments in WT and VE-cadherin-alpha-catenin chimera (VE-α-cat) mice (Figure 4D) showing no residual cells left in the bone marrow vessels, indicating that the detected cells were all homed into the bone marrow. Circles show occasional presence of cells in vessels. (D) Quantification of cell numbers present in the bone marrow vessels. No difference was detected. (E) Sixteen hours after administration of the VE-cadherin antibody, no permeability differences were detected anymore in both arterioles and sinusoids, indicting the reversible effects of blocking VE-cadherin using antibodies (n=2 per group).

**Supplemental figure 3:**
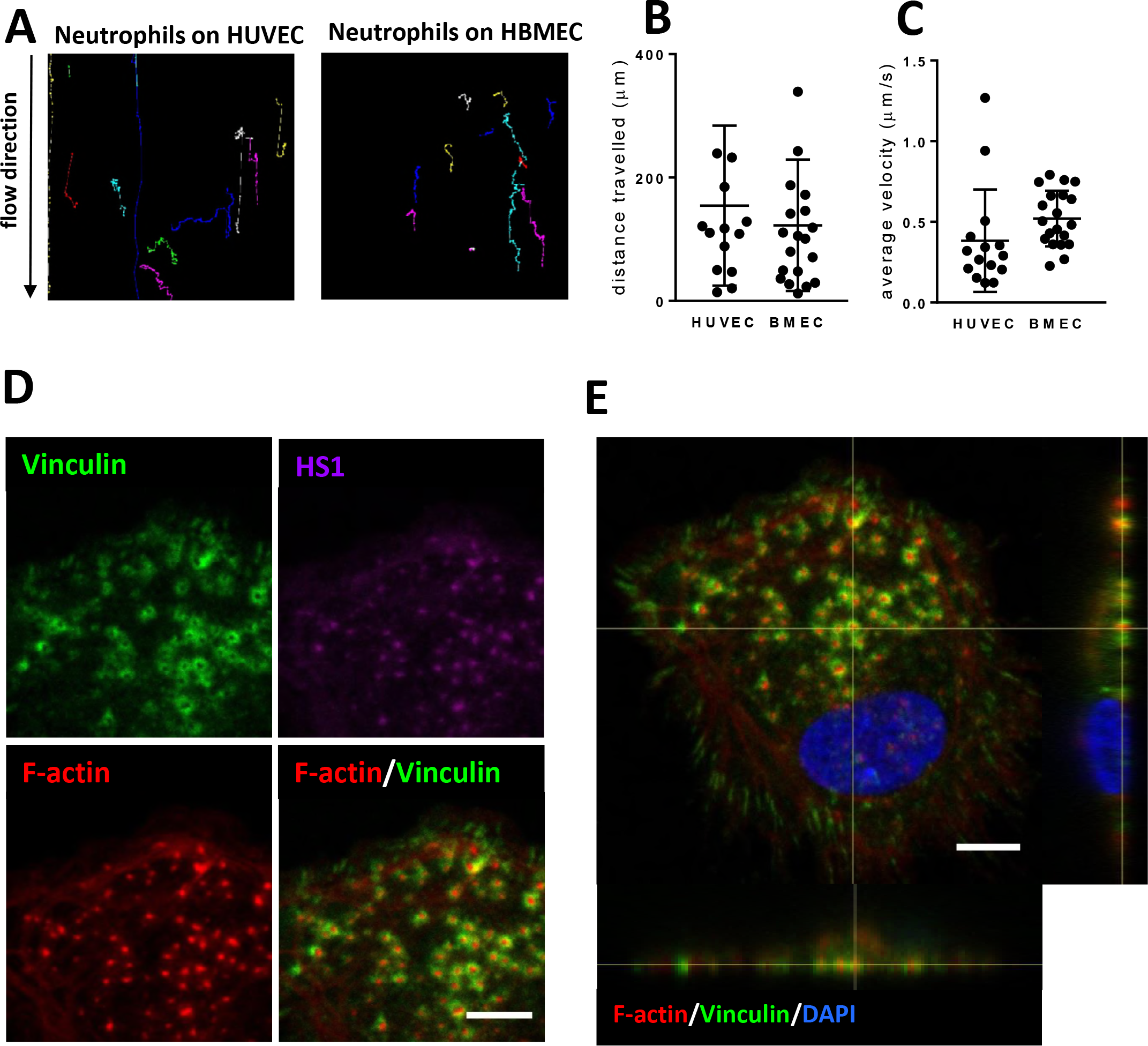
Migration of leukocytes over endothelium and podosome formation. (A) Migration tracks of cells under flow conditions on HUVEC (left) or HBMEC (right) show highly motile neutrophils. (B) Quantification of distance travelled on the apical surface of the endothelium in μm, or (C) average velocity (μm/s). Experiment was carried out independently at least three times. (D) Monocyte-derived dendritic cells were treated for 30 minutes with PMA showing the induction of podosomes, based on typical podosome markers: vinculin (green) in the circle and HS1 (magenta) and F-actin (red) in the core. (E) Orthogonal projection shows that podosomes localize at the basolateral surface with F-actin in red, vinculin in green and the nucleus in blue. Scale bar indicates 6μm.

